# InjectionDesign: Plate Design with Optimized Stratified Block Randomization for Modern LC/GC-MS-Based Sample Preparation

**DOI:** 10.1101/2023.02.26.530140

**Authors:** Miaoshan Lu, Hengxuan Jiang, Ruimin Wang, Shaowei An, Jiawei Wang, Changbin Yu

**Author notes:** Authors contributed equally (Miaoshan Lu, Hengxuan Jiang).

## Abstract

Plate Design is a necessary and time-consuming operation for GC/LC-MS based sample preparation. The implementation of the inter-batch balancing algorithm and the intra-batch randomization algorithm can have a significant impact on the final analysis results. For researchers without programming skills, a stable and efficient online service for plate design is necessary. However, most exist products do not currently have online services, and are not optimized for GC/LC-MS instruments with custom injection capabilities. Here we describe InjectionDesign, a free online plate design service focus on GC/LC-MS-based multi-omics experiment design. It offers the ability to separate the position design from the sequence design, making the output more compatible with the requirements of a modern mass spectrometer-based laboratory. In addition, it has implemented an optimized block randomization algorithm, which can be better applied to sample stratification with block randomization for unbalanced distribution. It is easy to use, with built-in support for common instrument models and quick export to worksheet. InjectionDesign is an open-source project based on Java. Researchers can get the source code of the project from Github: https://github.com/CSi-Studio/InjectionDesign. A free web service is also provided: http://www.injection.design.

**Demo Project:** http://www.injection.design/#/project/detail?projectId=Test

## 1. Introduction

Metabolomics or proteomics analysis based on mass spectrometry has great potential in scientific research. More laboratories have begun conducting metabolomics or proteomics analyses to effectively convert biological samples into digital samples. The factors that affect the final data quality, such as sample quality, experimental environment, instrument type, and sample pretreatment method, vary from laboratory to laboratory. The exploration and innovation of scientific research need to change some existing techniques. Still, the lack of some critical steps will undoubtedly lead to a decline in the quality of data. Exploring and understanding the experimental steps that need to be strictly followed requires much effort but is still worth doing [1–3]. Implementing quality assurance (QA) and quality control (QC) is an essential guarantee for the reliability and consistency of data[1]. The deviation of sample distribution will bring great interference to the computer experiment[2].

Randomizing the pretreated samples according to appropriate dimensions is an important method in the QC process for reducing machine drift and batch effect[4–6]. At present, there are already some tools for microplate design[7–9]. The software mentioned above is all developed in R language and only PlateDesigner provides web services. PlateDesigner can evenly distribute the samples according to a certain dimension, but only the completely random algorithm is used for the random algorithm in the plate. At the same time, PlateDesigner does not separate the sample position design from the injection sequence design, which does not apply to most modern mass spectrometers or chromatographs.

A large number of studies have shown that the batch effect of samples has a significant impact on the test results[5,10,11]. Among the many best practice recommendations, we focus on two standard and essential algorithms: inter-batch balancing and intra-batch randomization. This two-step can be easily manipulated by Excel or manually adjusted with simple randomization, but it becomes challenging when dealing with large queues and many participants, and a simple randomization algorithm would also introduce significant machine drift bias or batch effect bias. Operators will become prone to making mistakes under this circumstance. Different laboratories may use different models of LC-MS instruments to achieve optimal results. Different specifications of plates are also involved in the pretreatment process. More and more methods or standard materials for QC are being proposed[12]. However, there is less research on the information system used for the QC methods in the sample pretreatment stage, especially for the inter-batch balancing algorithm and intra-batch random algorithm, which is a recommended practice that has been proposed many times[2,13]. A stratified block randomization algorithm is an important and fast method to reduce the confounding variables’ interference[4].

With the increasing scale of studies, there are often thousands of samples in a single project. This project usually takes weeks or even months to complete which is unstable by generating random sequences at one time. In a subsequent implementation, the distribution needs to be adjusted and rearranged. Therefore, users hope that the sorting results can be traced and adjusted. Most modern high-throughput mass spectrometers have automatic sample injection procedures. Therefore, the position sequence of samples arranged on the plate does not indicate the sequence of sample injection. The instrument can customize the sampling sequence independently so that the layout of samples on the plate is more standardized and humanized. For example, we can put QC samples in the same area of the left part to reduce the operational complexity caused by QC sample interpenetration. This can greatly reduce the preprocessing time and error rate.

Here we describe InjectionDesign, a free web service focused on the sample pretreatment stage. InjectionDesign mainly solves the following problems:

1. Persistent storage capacity of sample injection information prepared for long-term projects.
2. Implementation of inter-batch balancing algorithm and intra-batch block randomization algorithm.
3. An optimized block randomization algorithm for unequally distributed samples.
4. Method for separating sample position layout from injection sequence design.

Appropriate sample pre-treatment methods are critical to the accuracy of mass spectrometry based data analysis. The randomization and balancing methods are rigorous and necessary. However, the introduction of a software for this purpose or the need for appropriate programming to obtain the appropriate work orders is cumbersome. An effective, free web service is very important to solve this problem. To meet the security requirements of some laboratories, we also provide additional downloadable local installation packages.

## 2. Materials and Methods

### 2.1. Inter-Batch Balancing Algorithm

The inter-batch balancing algorithm is suitable for the case of a known variable. To prevent system interference caused by aggregating features in the same batch. We must evenly distribute the samples with the same characteristics in each batch before injection, such as age, gender, and other factors. 81% of the laboratories reported the balancing step for samples[2].

The main detail of the inter-batch balancing algorithm lies in the treatment of the trailing samples. For example, we have a project with 50 males and 55 females. It needs to be stratified into 4 groups. After a simple equalization calculation, each group contains at least 12 males and 13 females. The tail sample is 2 males and 3 females. We randomized this tail of 5 samples again and interspersed them into 4 groups in turn.

### 2.2. Intra-Batch Randomization Algorithm

The block randomization algorithm combined with the stratified balancing algorithm is currently a commonly used random method in the sample pre-processing stage[4,5]. Its impact on subsequent data analysis has also been specifically discussed. The need to use block randomization algorithms is necessary in most projects[4]. The key to the block randomization algorithm is confirming the category amount of a given variable and setting the block size.

#### 2.2.1. For Uniformly Distributed Samples

Most biological experiments are well-designed before they begin. For example, there is a clear division between the control group and the observation group. At the same time, the number of samples in such groups is also evenly distributed, to better reduce the deviation of statistical results caused by the difference in sample distribution.

When we deal with an evenly distributed sample set, we have implemented a fast block randomization algorithm. For example, if we have a project of 100 samples, 50 are in the treatment group (samples in this group are defined as A), and 50 are in the placebo group (samples in this group are defined as B). As a result, the dim size is 2. Then the block size we can choose is 2, 4, 6, … We will use four as an example. Then the block permutation can be described as ABAB, BBAA, AABB, BABA, BAAB, and ABBA, a total of 6 combinations. Since each group contains 4 samples, a total of 25 groups are required. We need to generate a set of random number sequences in the range of 1-6. Then replace the combination of corresponding serial numbers into the sequence to obtain the final sequence. Figure.1(a),(b) shows the implementation principle of the block randomization algorithm.

**Figure.1.**
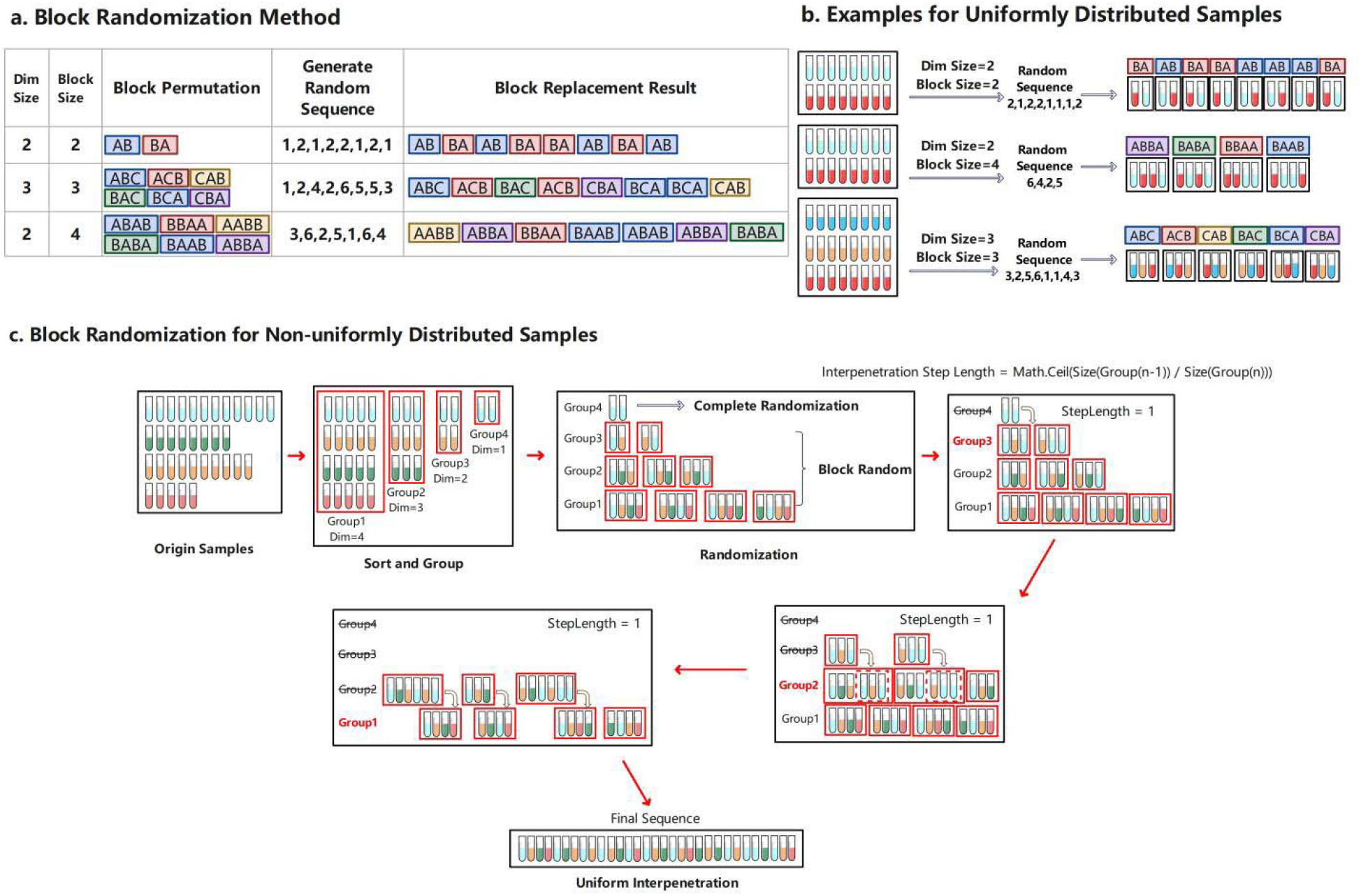
(a) The implementation principle of the block randomization algorithm. A block random factor with two variables needs to be confirmed: Dim size and block size. Then calculate the total number of block permutations according to the block random factor, calculate the number of blocks required according to the total number of samples and block size, and generate a corresponding number of random sequences. Finally, the corresponding combination is replaced in the randomized sequence to obtain the final distribution sequence. (b) The example random results under different block random factors, three factors (dim size is 2, block size is 2; dim size is 2, block size is 4; and dim size is 3, block size is 3) are shown here. (c) Optimized block randomization algorithm with group uniform interpolation method, which can maximize the uniformity of sample distribution after block randomization.

#### 2.2.2. For Non-uniformly Distributed Samples

Due to the scarcity of samples, many are unevenly distributed. This is also how most projects actually stand. We are unable to apply the block random approach directly when dealing with a queue that is not evenly distributed. This work presents an optimized block randomization approach for handling dispersed samples that are out of balance. See Figure.1(c). The unbalanced samples must first be sorted by the number of samples for each categorization, ascending. Sorted samples should be vertically grouped in accordance with the maximum dimension inclusion principle. The samples in each group are dispersed equally, and we can then obtain groups with various dimensions. Except for the one-dimension group that uses the complete randomization algorithm, all of the other groups use the block randomization algorithm because the sample distribution in each group is uniform. Following randomization, we then place the groups into adjacent groups. After that, by rounding down the length ratio of adjacent groups, we can determine the uniform step length for insertion. Then, in accordance with the step length, we combine with the corresponding blocks of the adjacent group to create new blocks with different lengths. The cycle is thus interspersed until the last two groups are merged.

### 2.3. Intra-Batch Position Design

The placement of the samples on the plate can be independent of the final injection sequence because the majority of contemporary mass spectrometers enable user-defined injection sequences. The effective delineation of the sample placement area on the plate can significantly increase sample placement efficiency and minimize placement errors.

Using QC samples to reduce batch effects and improve data quality is a common technical method in proteomics and metabolomics. InjectionDesign includes common QC sample types. Solvent QC Samples, Long Term Reference (LTR) QC Samples, Blank Samples, and so on. Each laboratory has its own QC plan based on the cost of the experiments. As a result, they will have a fairly fixed and specific QC layout plan. QC samples do not enter the instrument continuously in MS-based multi-omics research. According to certain rules, QC samples are generally evenly inserted into random samples.

See Figure.2. By placing QC samples in a specific area of the plate, Column A employs a horizontal or vertical layout that is suitable for human operation habits. This placement method simplifies not only the placement process but also the subsequent sample traceability. Column B is placed in the injection order, regardless of horizontal or vertical, and such QC layout will constantly be inconvenient. In laboratories without automated equipment, such a layout will result in a significant amount of extra work.

**Figure.2.**
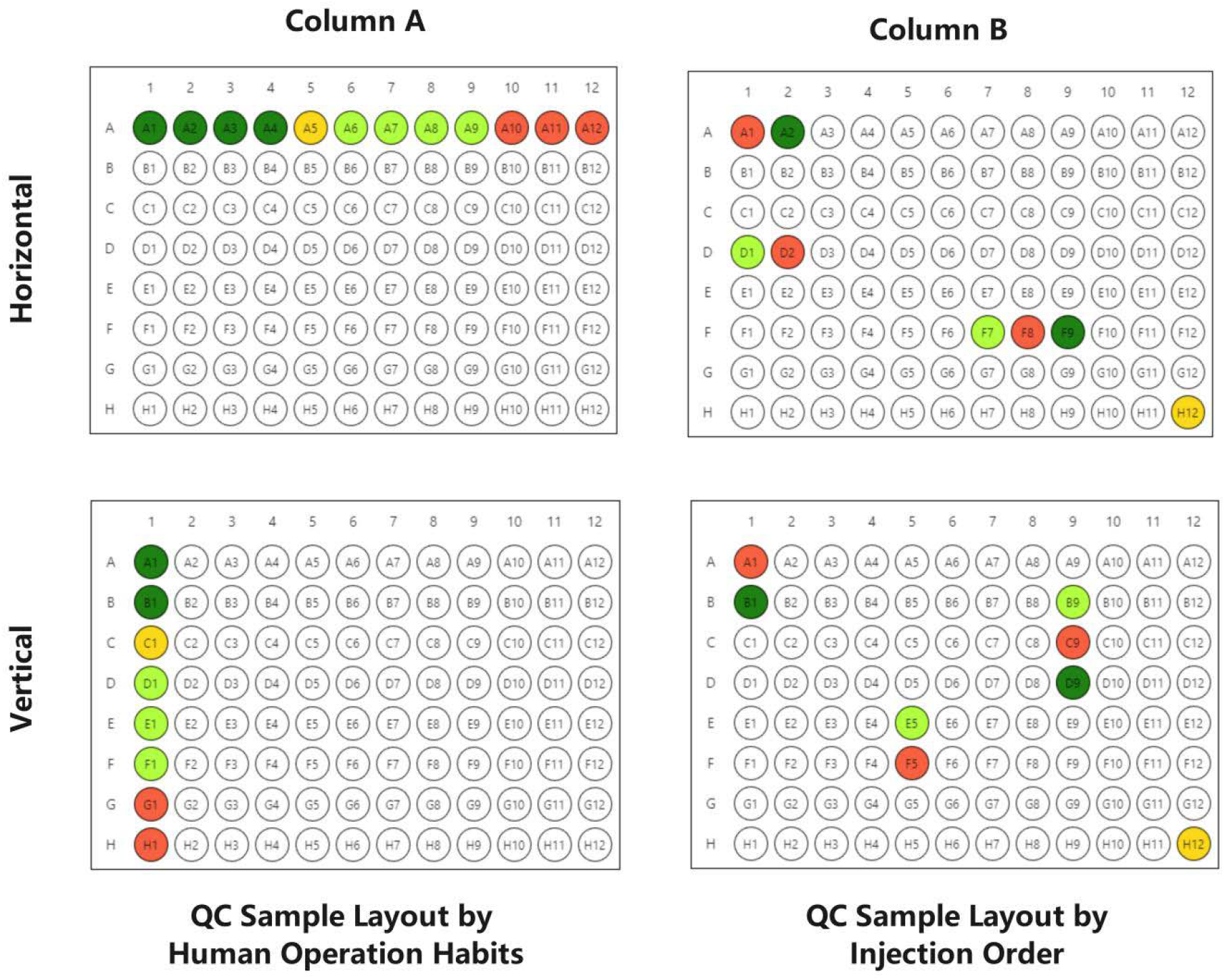
Visualization of QC samples in different layouts. It can be clearly seen that the ability to separate the plate position from the injection sequence brought by the customized injection sequence greatly improves the efficiency in sample pre-processing placement.

A complex process is involved in intra-batch position design. We can confirm the number of experimental samples processed in each batch after inter-batch balancing. This step must specify the location on the plate of each sample (including the QC samples) as well as the injection order. We built a workflow to make it simple to customize the steps for each laboratory. The step consists of four parts of information: Plate Specification. Plate Design Load. Predefined QC Location. Injection Order.

#### Plate Specification

Users must customize the plate’s specifications. The plate information includes x- and y-axis numbers as well as x- and y-axis labels. InjectionDesign includes the following plate templates: 96-well plate template (x-axis=12, x-labels=[1,2,3,4,5,6,7,8,9,10,11,12], y-axis=8, y-labels=[a,b,c,d,e,f,g,h]) and 384-well plate template.

#### Plate Load Capacity

Users should determine the number of QC samples of each type used. Select the custom plate’s maximum experimental sample capacity as well. We have built in many of the QC sample types suggested by mQACC[12], as well as custom sample types to allow for possible extensions for special samples in the lab.

#### Predefined QC Location

The predefined injection location is mainly for QC samples. QC samples are usually placed in the first or last column of the plate. Because there is no absolute relationship between the injection location and the injection order, it can be set according to the operator’s habits to reduce the difficulty of sample transfer between plates.

#### Pre-interpolation of QC

The QC pre-interpolation configuration is performed after randomization of the samples. QC samples are usually interpolated evenly among the experimental samples. It will randomly arrange all experimental samples before inserting QC samples in a circular fashion to form a new injection sequence.

## 3. Result

### 3.1 Main Workflow

A special workflow is formed as Figure.3. The user first needs to create a project to store all the imported sample information. Alternatively, samples can be imported directly via Excel. The system will automatically create the associated project. InjectionDesign provides the ability to store samples persistently. As a result, uploaded samples can be accessed and reprocessing repeatedly even after worksheets are generated.

**Figure.3.**
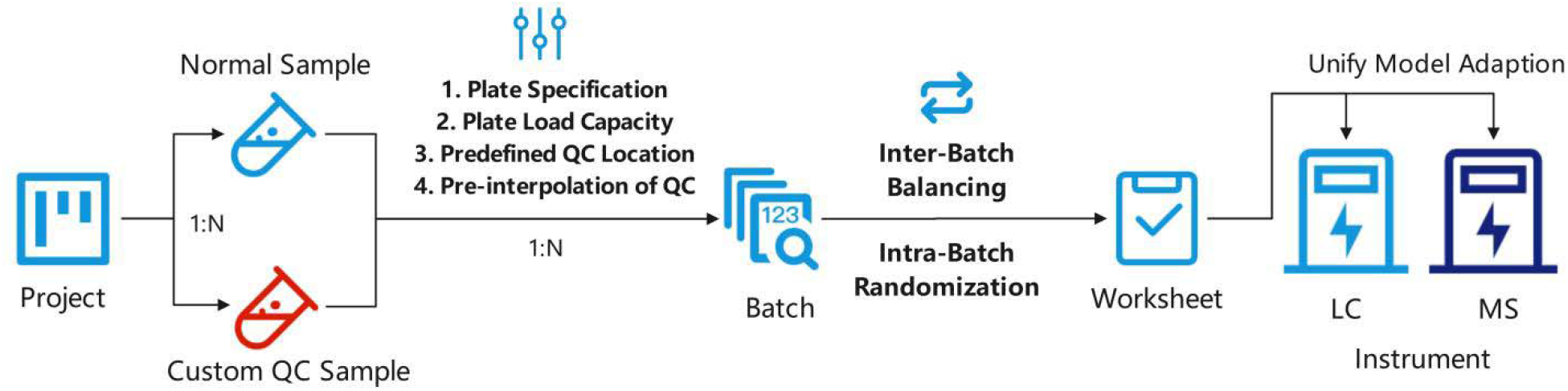
The main flow of InjectionDesign, which contains the implementation order of all the methods mentioned in the section.

### 3.2 Case Study

#### 3.2.1 Samples Preparation

We created a list of 1300 samples, each of which included the following information: gender (615 Male, 434 Female, 251 Unknown), whether treatment was used (650 treatment, 650 placebo), and age group (169 Kids, 351 Teenager, 481 Middle-Aged, 299 Older). See Figure.4a,4b. They represent two-category, three-category and four-category, respectively. The two-category is uniformly distributed. We chose a 96-well plate as the processing plate. The maximum number of samples that can be placed on each plate was also initially set to 96.

**Figure.4.**
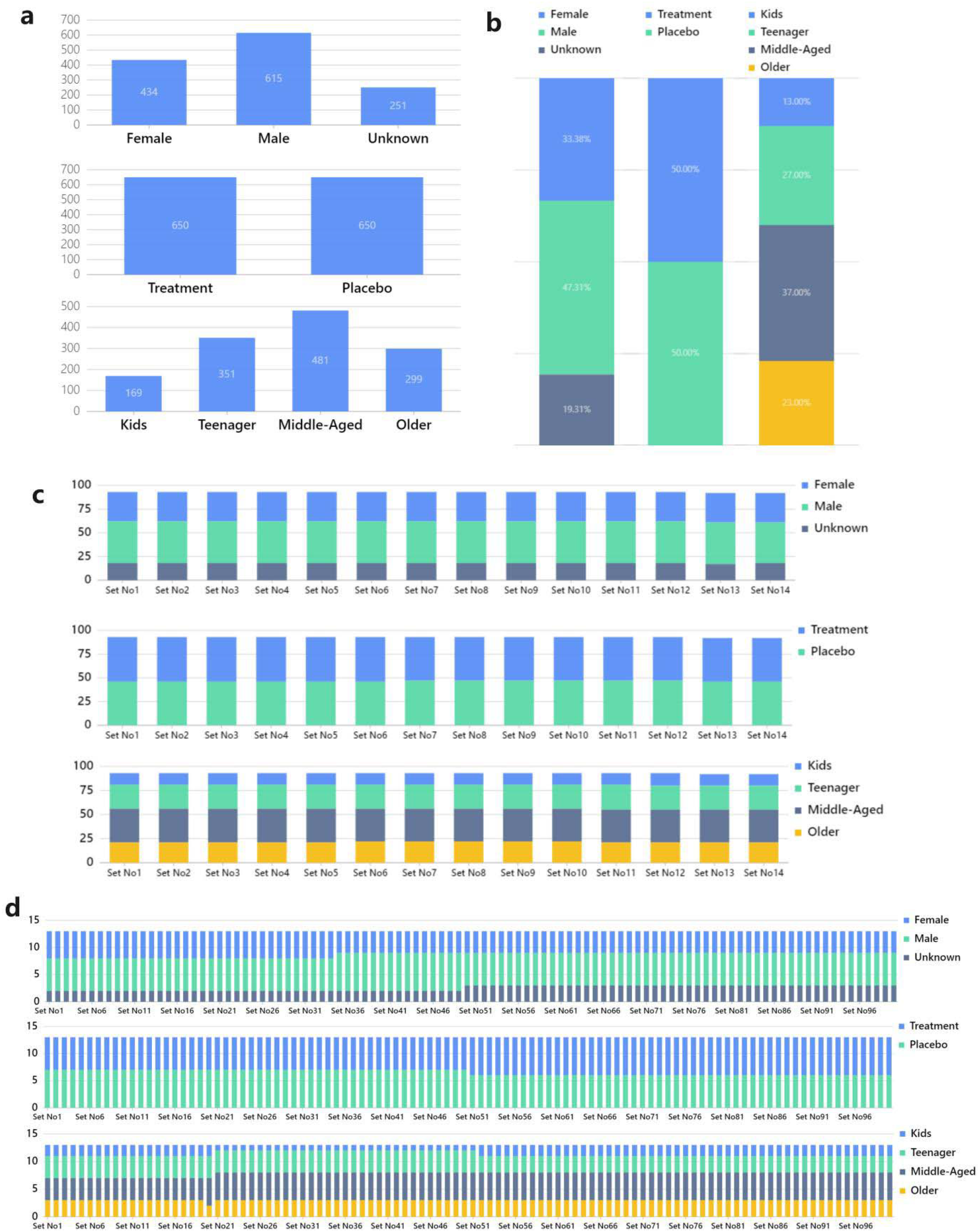
(a) Distribution of 1300 samples in three different dimensions. (b) Sample distribution in a stack bar viewer in three distinct dimensions. (c)(d) The proportion of samples distributed on each 96-well plate after using the inter-plate balancing algorithm. (c) specify the maximum number of samples on plate as 96, (d) specify the maximum number of samples on plate as 13. It is clear that, whether or not the distribution is balanced, the inter-plate balancing procedure works well for equating the sample partition.

#### 3.2.2 Inter-batch Balancing Result

Due to the enough sample number of individual plates, as seen in Figure.4c, the distribution of in single plate is extremely close to the overall ratio. To test the effectiveness of the balancing algorithm with a small number of samples. We set the maximum capacity of the plate to 13 (just for the convenience of division, set the number to about 10 can get the similar result). See Figure.4d. Although the very small number of individual plates leads to a significant imbalance in the proportion of unevenly distributed samples, the distribution of samples varies very little across the majority of the plate, but some plates do have slightly larger deviations. Naturally, we do not advise using a stratification sample size that is too small.

Additionally, InjectionDesign offers the complete randomization algorithm. This algorithm does not stratify by dimensions, it is completely randomly disrupted. This method works well for unidentified interferences or when simulating the sample’s actual distribution on a small scale.

#### 3.2.3 Intra-batch Randomization Result

The block randomization algorithm’s greatest benefit is that it guarantees that, in the event of a balanced distribution of samples, there will be absolutely no aggregated distribution. The treatment/placebo dimension was balanced, and since the block size is 2, theoretically there were no three consecutive samples of the same group in the results. See Figure.5a, In the final results, all samples were evenly distributed, and there were no three consecutive samples in the same group. In the remaining unbalanced dimensions, the optimized block randomization algorithm also achieves good results. All samples are relatively uniformly distributed, but still exhibits complete randomness in a small range.

**Figure.5.**
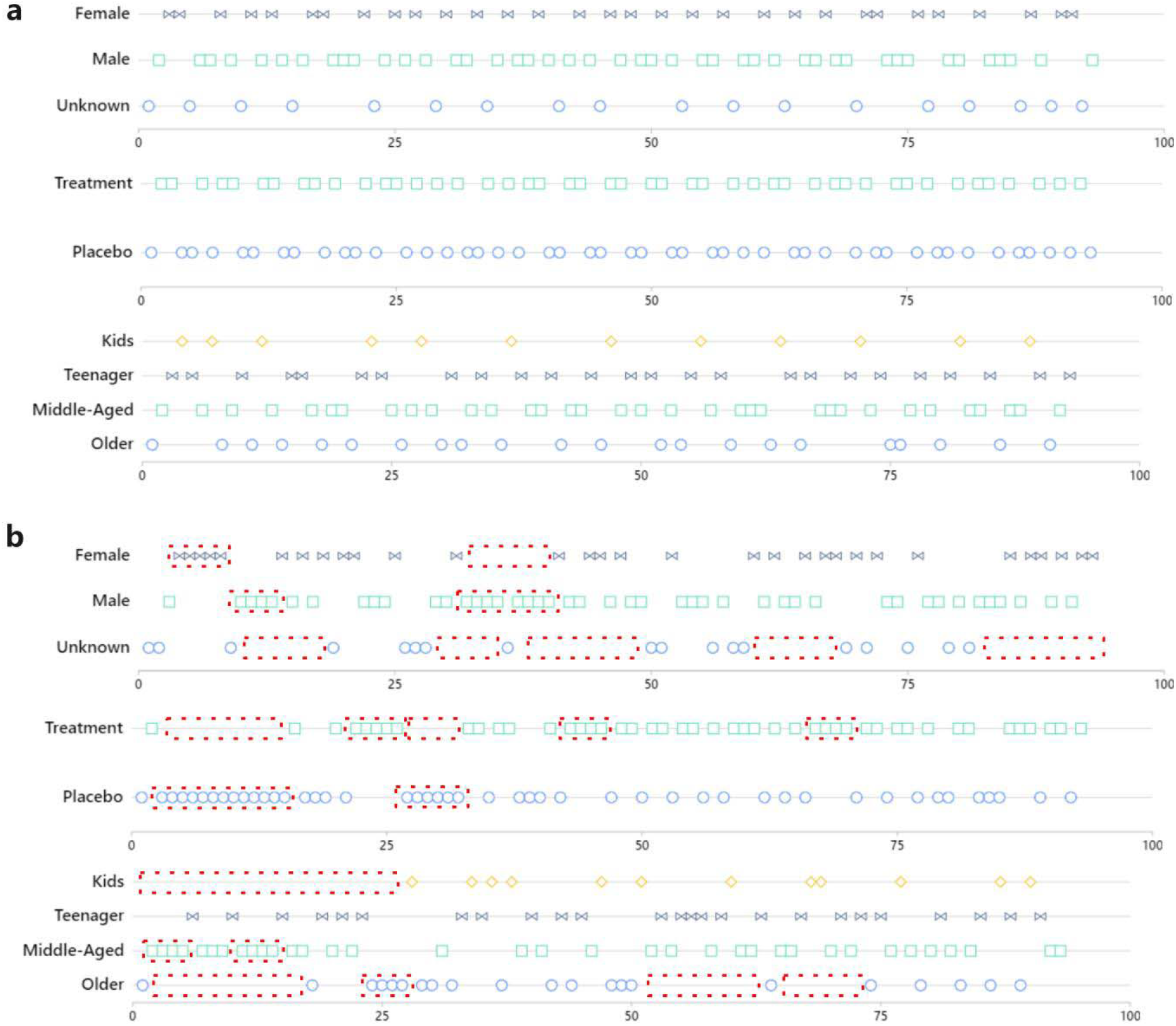
(a) Results of block randomization in three dimensions, the overall distribution of the data showed a clear balance. (b) Complete randomization result with three different dimensions. The area delineated by the red dashed box shows a clear random clustering of the sample distribution

Like the inter-batch balancing algorithm, InjectionDesign also provides a completely randomized algorithm for intra-batch samples. We also tested the same in three dimensions. The complete randomization algorithm generates a large number of aggregated features with high probability. See Figure.5b, the red box is the part with the more obvious aggregated distribution. The optimized block randomization algorithm has significant distribution control.

### 3.3 Real-time Visualization for Random Result

In the randomization step, users can adjust the parameters of stratified algorithm, randomization algorithm, stratified and randomization dimension. See Figure.6 After the parameters are set, the results of the randomized distribution are immediately displayed in a visual graph, which greatly helps the user to understand the final effect of the randomization.

**Figure.6.**
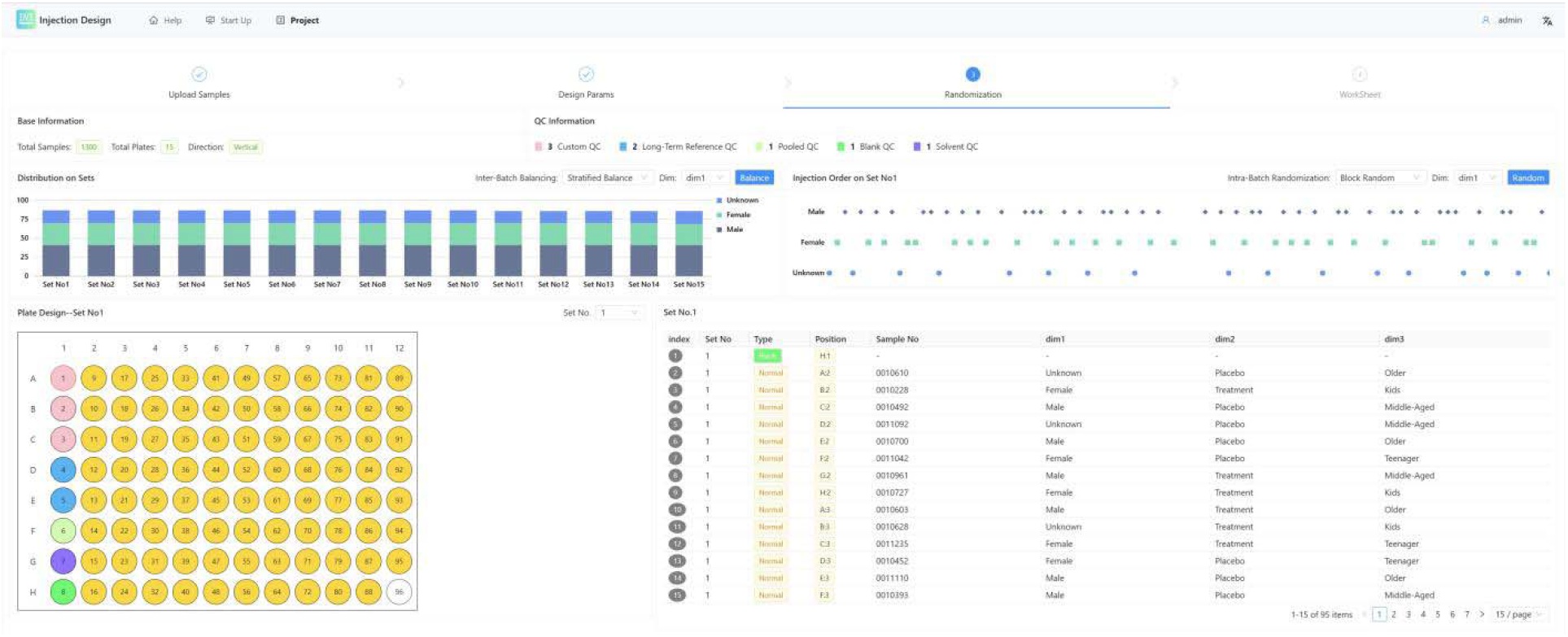
Screenshot of Injection for the randomiztion step.

## 4. Discussion

The sample processing flux in the laboratory is increasing as metabolomics and proteomics technology evolves. Some laboratories will install automated pre-processing equipment, but this is frequently costly. InjectionDesign focuses on the quality control steps of sample pretreatment, which can significantly reduce non-standard operation caused by manual processes. Despite the fact that many detailed steps are carried out in the form of SOP. Too many manual interventions, such as sample transfer between plates or sample injection tube label identification, will still result in errors. This will also be the source of power for InjectionDesign’s iterative process.

Stratified block randomization algorithm is suitable for most sample pre-processing. However, the block randomization algorithm completely averages out the distribution of the sample features, thus losing some of the numerical characteristics that are naturally present in the sample distribution. Although the complete randomization algorithm provided by InjectionDesign can be used to some extent for such scenarios, it is not a perfect solution.

Although the optimized block randomization algorithm can better balance the distribution of non-uniform samples. When there are too many groups in one dimension. Due to the averaging step calculation method, which can lead to local aggregation under some special numerical distributions. Better equilibrium results under extreme conditions may be achieved using solvers or related machine learning algorithms. InjectionDesign will also continue to update the kernel with new algorithms.

There are many established methods and procedures in current MS-based pre-treatment experiments. However, new methods and procedures are being developed all the time. For example, new QC samples and new injection rules may emerge. InjectionDesign also hopes to gain access to more of the lab’s innovative methodologies after the release, and to update them in the web service.

## 5. Conclusions

Sample pretreatment involves operators, samples, consumables, and instruments, the four most essential elements. This makes the operation complicated and prone to human errors. InjectionDesign provides various laboratories with rich extensibility by customizing QC types, randomization methods, plate types, export fields and injection rules. To enable each laboratory to try InjectionDesign quickly, we provide a free web service with suitable visual interfaces.

Because of the data security requirement in some laboratories, InjectionDesign also provides the ability to install locally. InjectionDesign provides a complete installation package on Github. InjectionDesign relies on community MongoDB and OpenJDK and is a cross-platform deployable application. Users can also make suggestions directly on GitHub, and the development team will respond quickly.

Author Contributions: Methodology, Miaoshan Lu and Changbin Yu; Software, Hengxuan Jiang, Ruiming Wang, Shaowei An and Jiawei Wang; Writing – review & editing, Ruiming Wang and Shaowei An. All the authors have revised and approved the final manuscript. All authors have read and agreed to the published version of the manuscript.

## Institutional Review Board Statement

Not applicable.

## Informed Consent Statement

Not applicable.

## Data Availability Statement

Code is found at https://github.com/CSi-Studio/InjectionDesign.

## Abbreviations

The following abbreviations are used in this manuscript:

LC: Liquid Chromatograph
GC: Gas Chromatograph
QC: Quality Control
QA: Quality Assurance
MS: Mass Spectrometry
mQACC: Metabolomics Quality Assurance and Quality Control Consortium
LTR: Long-Term Reference

## References

1. Beger, R.D.; Dunn, W.B.; Bandukwala, A.; Bethan, B.; Broadhurst, D.; Clish, C.B.; Dasari, S.; Derr, L.; Evans, A.; Fischer, S.; et al. Towards Quality Assurance and Quality Control in Untargeted Metabolomics Studies. Metabolomics 2019, 15, doi:10.1007/s11306-018-1460-7.

2. Evans, A.M.; O’Donovan, C.; Playdon, M.; Beecher, C.; Beger, R.D.; Bowden, J.A.; Broadhurst, D.; Clish, C.B.; Dasari, S.; Dunn, W.B.; et al. Dissemination and Analysis of the Quality Assurance (QA) and Quality Control (QC) Practices of LC–MS Based Untargeted Metabolomics Practitioners. Metabolomics 2020, 16, doi:10.1007/s11306-020-01728-5.

3. Bouhifd, M.; Beger, R.; Flynn, T.; Guo, L.; Harris, G.; Hogberg, H.; Kaddurah-Daouk, R.; Kamp, H.; Kleensang, A.; Maertens, A.; et al. Quality Assurance of Metabolomics. ALTEX 2015, 32, doi:10.14573/altex.1509161.

4. Burger, B.; Vaudel, M.; Barsnes, H. Importance of Block Randomization When Designing Proteomics Experiments. J Proteome Res 2021, 20.

5. Bailey, R.A. Design of Comparative Experiments; 2008;

6. Surowiec, I.; Johansson, E.; Stenlund, H.; Rantapää-Dahlqvist, S.; Bergström, S.; Normark, J.; Trygg, J. Quantification of Run Order Effect on Chromatography - Mass Spectrometry Profiling Data. J Chromatogr A 2018, 1568, doi:10.1016/j.chroma.2018.07.019.

7. Yan, L.; Ma, C.; Wang, D.; Hu, Q.; Qin, M.; Conroy, J.M.; Sucheston, L.E.; Ambrosone, C.B.; Johnson, C.S.; Wang, J.; et al. OSAT: A Tool for Sample-to-Batch Allocations in Genomics Experiments. BMC Genomics 2012, 13, doi:10.1186/1471-2164-13-689.

8. Suprun, M.; Suárez-Fariñas, M. PlateDesigner: A Web-Based Application for the Design of Microplate Experiments. Bioinformatics 2019, 35, doi:10.1093/bioinformatics/bty853.

9. Borges, H.; Hesse, A.M.; Kraut, A.; Couté, Y.; Brun, V.; Burger, T. Well Plate Maker: A User-Friendly Randomized Block Design Application to Limit Batch Effects in Large-Scale Biomedical Studies. Bioinformatics 2021, 37, doi:10.1093/bioinformatics/btab065.

10. Zhang, T.; Gaffrey, M.J.; Monroe, M.E.; Thomas, D.G.; Weitz, K.K.; Piehowski, P.D.; Petyuk, V.A.; Moore, R.J.; Thrall, B.D.; Qian, W.J. Block Design with Common Reference Samples Enables Robust Large-Scale Label-Free Quantitative Proteome Profiling. J Proteome Res 2020, 19, doi:10.1021/acs.jproteome.0c00310.

11. Statistical Analysis of Proteomics, Metabolomics, and Lipidomics Data Using Mass Spectrometry; 2017;

12. Lippa, K.A.; Aristizabal-Henao, J.J.; Beger, R.D.; Bowden, J.A.; Broeckling, C.; Beecher, C.; Clay Davis, W.; Dunn, W.B.; Flores, R.; Goodacre, R.; et al. Reference Materials for MS-Based Untargeted Metabolomics and Lipidomics: A Review by the Metabolomics Quality Assurance and Quality Control Consortium (MQACC), 2022, Vol. 18.

13. Broadhurst, D.; Goodacre, R.; Reinke, S.N.; Kuligowski, J.; Wilson, I.D.; Lewis, M.R.; Dunn, W.B. Guidelines and Considerations for the Use of System Suitability and Quality Control Samples in Mass Spectrometry Assays Applied in Untargeted Clinical Metabolomic Studies. Metabolomics 2018, 14.

